# Spleen-dependent role of cyclooxygenase-1 in the physiological manifestations of severity in systemic inflammation

**DOI:** 10.64898/2026.07.07.737102

**Authors:** Camila F. Brito, Eduardo H. Moretti, Isis F. L. Trzan, Monique T. Fonseca, Lucas M. M. Marques, Jady T. Guedes, Evilin N. Komegae, Elizabeth A. Flatow, Norberto P. Lopes, Alexandre A. Steiner

## Abstract

Cyclooxygenase-1 (COX-1) is classically regarded as a constitutive enzyme that produces eicosanoids with housekeeping functions, but recent evidence indicates that it may also be involved in the acute phase of severe systemic inflammation. There is evidence indicating that COX-1 is selectively activated in the spleen via post-translational mechanisms early the course of LPS-induced systemic inflammation. However, the mechanistic link between COX-1 and the spleen has not yet been demonstrated in direct experiments. The present study was conducted to fill this gap. The effects of the COX-1 inhibitor SC-560 on the LPS-induced severity triad (hypotension, hypothermia and acidosis) were evaluated in rats subjected to splenectomy or in sham-operated controls. In the sham-operated group, SC-560 significantly attenuated the severity triad independently of changes in plasma cytokines (TNF and IL-1β). In the splenectomized rats, SC-560 completely lost its ability to attenuate the hypotension and the acidosis induced by LPS. The effect of SC-560 on LPS-induced hypothermia was also impaired by splenectomy, though not completely. We then conducted a lipidomic screening to identify which COX-1-derived eicosanoids might be responsible for mediating the severity triad. Based on spleen-blood correlations, the screening identified PGE_2_ and PGD_2_ as putative candidates. In conclusion, the present study provides direct evidence for a mechanistic link between the spleen and COX-1 in the mediation of severity in systemic inflammation, and identifies PGE_2_ and PGD_2_ as putative candidates involved.

## 1. Introduction

Cyclooxygenase (COX) — *a.k.a.*, prostaglandin H synthase — is a rate-limiting step in the biosynthesis of eicosanoids (Flower, 2019; Mitchell & Kirkby, 2019). In the 1990s, the discovery that there are two COX isoforms set the foundation for the concept that the COX-1 isoform is constitutively expressed and plays housekeeping roles, whereas the COX-2 isoform is inducible and accounts for the inflammation-related roles of eicosanoids (Simmons *et al*., 2004; Dennis & Norris, 2015). This dichotomy profoundly influenced the pharmacology of non-steroidal anti-inflammatory drugs, driving the development of selective COX-2 inhibitors as a strategy to achieve anti-inflammatory and analgesic effects while mitigating the gastrointestinal adverse effects of this drug class (Simmons *et al*., 2004; Dennis & Norris, 2015).

More recently, however, this dichotomous model has proven to be an oversimplification, as some studies have uncovered that COX-1 is not devoid of roles in inflammation under certain contexts. In a murine model of flagellin-induced severe systemic inflammation, an early COX-1-dependent eicosanoid storm has been shown to be an early driver of mortality (von Moltke *et al*., 2012). Such a response was driven by NLRC4 inflammasome in a cytokine-independent fashion (von Moltke *et al*., 2012). By the same token, selective COX-1 inhibition in a rat model of systemic inflammation induced by bacterial lipopolysaccharide (LPS) has been found to prevent the development of hypotension and hypothermia (Akarsu & Mamuk, 2007; Steiner *et al*., 2009), which, along with metabolic acidosis (Steiner *et al*., 2017), form a triad of physiological alterations typically associated with severe forms of systemic inflammation. Together, these lines of evidence indicate that while COX-1 is not involved in localized inflammatory processes or less severe forms of systemic inflammation, it plays pivotal roles in the early stages of severe inflammatory responses.

In a previous study (Steiner *et al*., 2009), the enzymatic activity of COX-1 has been shown to be heightened in the spleen of LPS-challenged rats, but not in the liver, lungs, kidneys or brain. Such activation involved post-translational mechanisms, since neither COX-1 mRNA nor protein was altered in the spleen. However, the mechanistic link between COX-1 and the spleen has not yet been demonstrated in direct experiments. The present study was conducted with the goal of filling out this important gap. The severity of systemic inflammation was assessed based in a triad of physiological manifestations: hypotension, hypothermia and acidosis. The latter of these manifestations was inferred by an acute rise in the respiratory exchange ratio (RER), as previously described (Steiner *et al*., 2017). COX-1 was inhibited pharmacologically using SC-560 (Smith *et al*., 1998). To test whether the spleen is, in fact, the source of the COX-1-derived signals, experiments involving a combination of splenectomy and COX-1 inhibition were performed. Cytokines were measured, and a targeted lipidomics approach was employed to screen for the putative mediators of the COX-1-dependent effects.

## 2. Materials and Methods

### 2.1. Animals

The study was conducted in male Wistar rats aged 8-12 weeks. The rats were obtained from the specific pathogen-free facility of the Universidade de Sao Paulo. They were housed at an ambient temperature of 24-26°C and under a 12:12 hour light-dark cycle (lights on at 7 AM). Standard rodent chow and water were available *ad libitum*. The rats were housed in groups of three per cage before the surgical preparation and individually after surgery. All protocols were approved by the Institutional Animal Care and Use Committee.

### 2.2. Drugs

LPS from *E. coli* O55:B5 (Sigma-Aldrich) was delivered intravenously as a 1-mg/ml suspension in saline, in a volume of 1 ml/kg, resulting in a dose of 1 mg/kg. The injection was performed via a pre-implanted venous (jugular) catheter (see *Surgical Preparation* and *Experimental Setup* below). To break up self-aggregates of LPS, the suspension was sonicated in bath for 30 min immediately before use.

The COX-1 inhibitor, SC-560 (Cayman Chemical), was administered by an intraperitoneal injection, performed 90 min before the LPS injection. SC-560 was dissolved in a vehicle consisting of propylene glycol (30%), ethanol (30%) and water (40%). The concentration of SC-560 in the solution was 5 mg/ml, which was administered in a volume of 1 ml/kg, providing the dose of 5 mg/kg.

### 2.3. Surgical preparation

In preparation for an experiment, one or more of the following surgical procedures was performed: implantation of a pressure/temperature telemetry transmitter; implantation of an intravenous (jugular) catheter; and total splenectomy. Antibiotic prophylaxis with enrofloxacin (5 mg/kg s.c.) was provided immediately before surgery. The surgeries were performed aseptically under anesthesia with isoflurane (1.5-2.5%). Body core temperature was maintained at 36-37°C with the help of an isothermal pad.

For implantation of the telemetry transmitter, a midline laparotomy was performed. The telemetry transmitter (model C50-PT; Data Sciences International) consisted of a capsule joined to a gel-filled catheter, which was implanted nonocclusively in the abdominal aorta. The abdominal aorta was accessed by displacing the small intestine sideways with saline-soaked gauze pads. The aorta was temporarily ligated caudal to the renal arteries while the catheter was being inserted and secured in place with octyl-cyanoacrylate glue. Blood flow was restored within 2 min. The capsule of the pressure transmitter was sutured to the ventral muscles during the closing process.

For venous catheterization, the left external jugular was isolated and permanently ligated. A 3-Fr polyurethane catheter was inserted into the vein caudally to the ligation site, and then advanced until its tip reached the right atrium. The distal end of the catheter was tunneled under the skin and exteriorized at the nape, after which the catheter was locked with heparinized glycerol (500 U/ml).

Splenectomy was performed via a dorsoventral incision made near the left costal margin of the thorax, as previously described (Fonseca *et al*., 2021). Briefly, the spleen was pulled through the incision, the splenic vessels were ligated, and the organ was excised.

The rats were allowed to recover from surgery for one week before an experiment. For pain management, the rats received ketoprofen (5 mg/kg s.c.) at the end of surgery and once a day in the following two days.

### 2.4. Experimental Setup

The experiments were conducted one week after the surgical preparation. On the day of the experiment, unanesthetized, freely moving rats were transferred to an environmental chamber (Environmental Growth Chambers) set to 22°C, an ambient temperature that is preferred by rats in the early stage of LPS-induced hypotension (Almeida *et al*., 2006). Inside chamber, each rat was housed inside a 5 L open-flow respirometry cage. The cage was sealed, except for a 2.5 cm opening at the top for air intake and infusion harness/swivel system pass-through, as well as for three 4-mm hose connectors on the sides, from where air was pulled at a rate of ∼1800 ml/min with the help of a SS4 subsampler unit (Sable Systems). The air entering and exiting the respirometry cage was sampled at 5 min intervals, and the fraction of oxygen in each sample was determined by a FOXBOX gas analyser (Sable Systems). The analog outputs of the SS4 unit and of the FOXBOX gas analyser were converted to digital by the UI-2 data acquisition interface, and acquired in a computer using the Expedata software (Sable Systems). The rate of oxygen consumption (VO2) and the respiratory exchange ratio (RER) were calculated as described in *Data Processing and Analyses*.

Inside the respirometry cage, each rat was equipped with an infusion harness, which protected a PE-50 extension of the i.v. catheter. The catheter extensions were passed by a swivel system (Instech) and connected to a syringe located outside the environmental chamber, from where i.v. injections and infusions were performed. For the i.p. injection, the respirometry cage had to be temporarily open, but this took place 90 min before the LPS injection. A PhysiolTel RPC-1 receiver (Data Sciences) was positioned under each cage; it captured the radio waves emitted by the telemetry transmitter. The signals captured by the telemetry receiver were conveyed to a computer and recorded at 500 Hz with the help of the Data Sciences ART software.

### 2.5. Tissue harvesting

At the time corresponding to the maxima of hypothermia, hypotension and acidosis (80 min post-LPS), the rats were anesthetized with thiopental (10 mg/kg) administered via the i.v. catheter extension. A drop of blood was collected from the animal’s tail and used for determination of clotting time by the capillary tube method; clotting time was used to confirm the effectiveness of COX-1 inhibition by SC-560. The thoracic cavity was then opened, so that a larger amount of blood could be collected from the inferior vena cava into heparinized tubes for lipidomic analysis and for determination of cytokine concentrations. When present (non-splenectomized rats), the spleen was also harvested for lipidomic analysis; it was cleared of blood prior to harvesting by transcardial perfusion with cold phosphate-buffered saline.

### 2.6. LC-MS/MS-based lipidomics

Plasma was obtained by centrifugation (4,500 × *g*, 5 min, 4°C) of the heparinized blood samples. Spleen extracts were prepared by: (i) homogenization in ice-cold methanol containing indomethacin and BHT; (ii) removal of insoluble remains by centrifugation (4,500 x *g*, 4°C, 10 min); (iii) evaporation of methanol under a stream of nitrogen at room temperature; and (iv) dissolution of the residue in water.

After the internal standard (PGE_1_) was added, the samples were acidified with 0.1% formic acid. The samples were then subjected to solid-phase extraction in an Oasis mixed-mode anion exchange μElution plate (Waters) in three sequential steps: (i) The plate cartridge was preconditioned with methanol and then with water; (ii) samples were loaded into the cartridge, and unwanted compounds were eluted in water containing 5% of methanol; and (iii) the compounds of interest were eluted in methanol containing 0.1% of formic acid. The prepared samples were then passed through an Acquity UPLC system equipped with a BEH C18 column (1.7-μm particles, 2.1 mm by 50 mm), which was coupled to an Acquity TQD MS/MS detector. The mobile phase consisted of water (solvent A) and acetonitrile (solvent B), both with 0.1% formic acid, and it was pumped at a flow rate of 0.4 ml/min. The gradient elution program was performed in four stages: (i) solvent B at 35% for 6 min, (ii) solvent B at 90% for 1.5 min, (iii) solvent B at 99% for 2.5 min, and (iv) solvent B at 30% for 3 min. The injection mode was full loop, with injection volume of 10 μl. The MS tune parameters were as follows: 0.16 liter/hour for the flow of the collision gas (argon); 3.02 × 10^−3^ mbar for the pressure in the collision cell; and 1,000 liters/hour for the flow of the desolvation gas (N2). The source and desolvation temperatures were 150° and 400°C, respectively. A multiple reaction monitoring mode was used for the quantification of the analytes in negative ionization mode, using the parameters described elsewhere (Fonseca *et al*., 2021).

Primary prostanoids and their stable metabolites were evaluated using standards purchased from Cayman: thromboxane B_2_ (TXB_2_, a TXA_2_ metabolite); prostaglandin F_2α_ (PGF_2α_, primary prostanoid); 6-keto Prostaglandin F1α (k-PGF1α; a PGI_2_ metabolite); 15-deoxy-Δ^12,14^-prostaglandin J_2_ (do-PGJ_2_, a PGD_2_ metabolite); tetranor-prostaglandin D metabolite (t-PGDM); prostaglandin D_2_ (PGD_2_, primary prostanoid); 13,14-dihydro-15-keto-prostaglandin E_2_ (d,k-PGE_2_, PGE_2_ metabolite); and prostaglandin E_2_ (PGE_2_, primary prostanoid).

### 2.7. Cytokine assay

Plasma was obtained by centrifugation (4,500 × *g*, 5 min, 4°C) of the heparinized blood samples. The concentrations of TNF, IL1β, IL6 and IL10 in the plasma samples were determined by sandwich ELISA using reagents and protocols from R&D Systems.

### 2.8. Data processing and analysis

Continuous recordings of arterial pressure (AP) and core body temperature (T_c_) were processed and analysed in the Ponemah software (Data Sciences). The open-flow respirometry data were initially processed in Expedata, then subjected to calculations of the following equations in MS Excel:

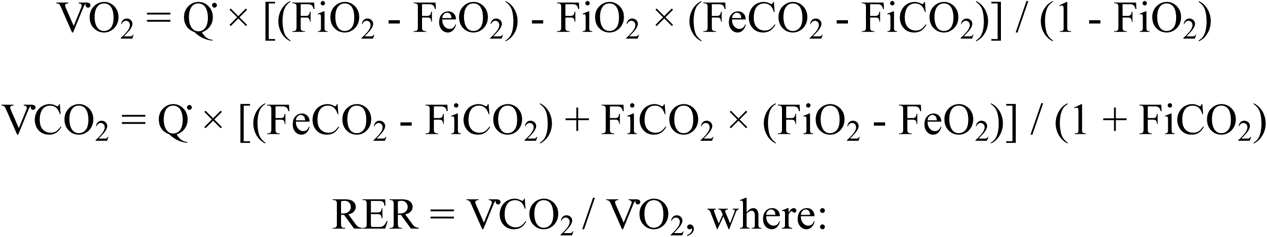

VO_2_ is the rate of oxygen consumption, VO_2_ is the rate of CO_2_ production; RER is the respiratory exchange ratio; Q is the air flow; FiO_2_ is the inspired fraction of O_2_; FeO_2_ is the expired fraction of O_2_; FiCO_2_ is the inspired fraction of CO_2_; and FeCO_2_ is the expired fraction of CO_2_.

Mean AP (AP_mean_), T_c_ and RER were plotted as change (Δ) from baseline (averaged over the 30 min that preceded the LPS challenge). Eicosanoid concentrations in the plasma and spleen were expressed in relative terms, using the internal standard as reference. Intergroup (SC-560 to vehicle) differences in eicosanoid concentration were compared across plasma and spleen; an equivalence plasma-spleen line was traced and used as reference. Residuals from the equivalence line were calculated for each analyte. TNF and IL1β concentrations are shown in absolute values (ng/ml).

Statistical comparisons were performed using Statistica 8.0 (StatSoft) and SAS Enterprise 9.2, with the level of significance set to 0.05. Baseline values of AP_mean_, T_c_ and RER were compared by one-way ANOVA. The time-dependent (post-LPS) changes in these parameters were evaluated using a linear mixed-effects model, with treatment (SC-560 *vs.* vehicle) set as the fixed effect, time as the repeated measurements and subjects as the random effect. Clotting time, as well as the concentrations of eicosanoids and cytokines were evaluated for the effect of treatment (SC-560 *vs.* vehicle) using Student’s *t* test.

## 3. Results

The biological effectiveness of SC-560 was confirmed based on its known influence on blood clotting time: 190 ± 23 s (mean ± SEM) in the SC-560-pretreated group *versus* 145 ± 15 s in the vehicle-pretreated group (*p* < 0.05). In the presence of the spleen (sham-operated rats), SC-560 attenuated by the same magnitude the LPS-induced decreases in both systolic and diastolic AP (data not shown), in this way attenuating the fall in AP_mean_ (Fig. 1A). In the same time window (15-180 min post-LPS), SC-560 significantly attenuated the hypothermic manifestation of systemic inflammation (Fig. 1B), as well as the biphasic rise in RER (Fig. 1C). The early rise in RER during LPS-induced systemic inflammation has been validated as a proxy of metabolic acidosis (Steiner *et al*., 2017), and was used herein for having the advantage of being recorded noninvasively for the duration of the experiment, which cannot be accomplished by blood gasometry.

**Figure 1.**
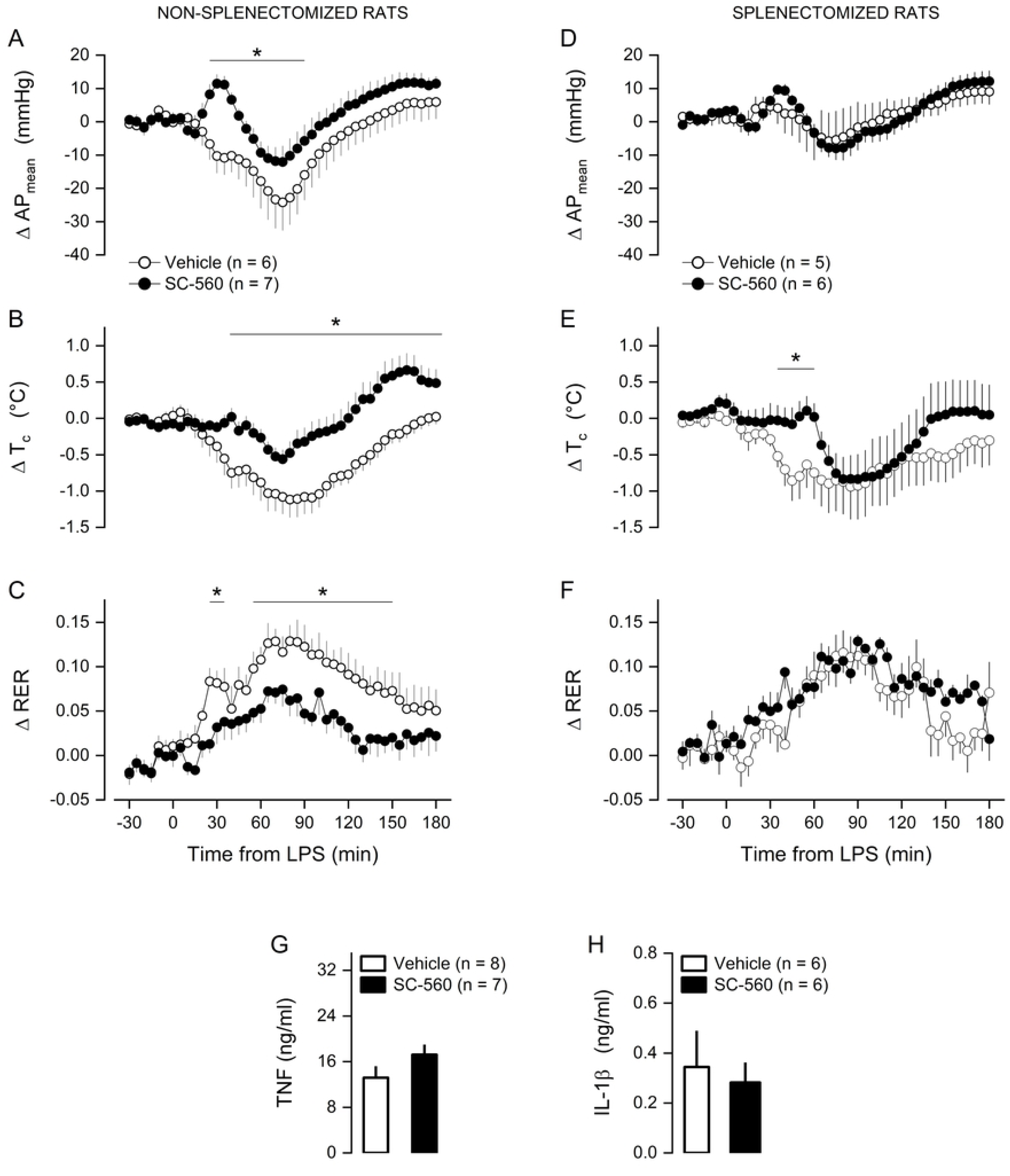
Effects of SC-560 on cardiovascular and thermometabolic parameters of the LPS-challenged rats subjected to splenectomy or to sham operation (non-splenectomized). Mean arterial pressure (AP_mean_), core body temperature (T_c_) and respiratory exchange ratio (RER) are shown in panels **A-C** for the non-splenectomized rats and in panels **D-F** for the splenectomized rats. An acute rise in RER in this model has been shown to be a proxy of metabolic acidosis in the rat model of LPS-induced systemic inflammation (see text for details). Plasma concentrations of TNF and IL-1β at 80 min post-LPS are shown in panels **G-H**. SC-560 or its vehicle was injected 90 min before the LPS challenge. Data are presented as change (Δ) from the baseline period, defined as the 30 min that preceded the LPS injection. Values are mean ± SEM. The number of animals per group (n) is indicated. *Statistically significant difference between the SC-560 and vehicle groups. There was no statistical difference in baseline values. Baseline values for APmean were: 116 ± 9 mmHg for the vehicle-pretreated, non-splenectomized rats; 110 ± 9 mmHg for the vehicle-pretreated, splenectomized rats; 110 ± 6 mmHg for the SC-560-pretreated, non-splenectomized rats; 116 ± 15 mmHg for the SC-560-pretreated, splenectomized rats. Baseline values for Tc were: 37.4 ± 0.5°C for the vehicle-pretreated, non-splenectomized rats; 37.3 ± 0.6°C for the vehicle-pretreated, splenectomized rats; 37.3 ± 0.4°C for the SC-560-pretreated, non-splenectomized rats; 37.1 ± 0.5°C for the SC-560-pretreated, splenectomized rats. Baseline values for RER were: 0.90 ± 0.06 for the vehicle-pretreated, non-splenectomized rats; 0.89 ± 0.05 °C for the vehicle-pretreated, splenectomized rats; 0.89 ± 0.05 °C for the SC-560-pretreated, non-splenectomized rats; 0.90 ± 0.05 °C for the SC-560-pretreated, splenectomized rats.

Next, we asked whether the spleen could be the site from which COX-1 exerts such effects. To test this hypothesis, we evaluated the effects of SC-560 on the LPS-induced responses in rats that had been subjected to surgical removal of the spleen. In the splenectomized rats, SC-560 lost its ability to attenuate the hypotensive and the RER (acidosis) responses induced by LPS (Figs. 1D and 1F). The effect of SC-560 on LPS-induced hypothermia was also impacted by splenectomy, but not completely. More specifically, in the absence of the spleen, SC-560 was still able to prevent the development of hypothermia till 60 min post-LPS, but lost its ability to impact this response after that (Fig. 1E). Taken together, these findings provide direct evidence for the existence of COX-1-dependent mechanisms in the spleen that mediate the severity triad in the early stages of endotoxic shock.

Because pro-inflammatory cytokines are known to mediate hypotension, hypothermia and acidosis in systemic inflammation (Tracey *et al*., 1986; Boomer *et al*., 2011; Fonseca *et al*., 2016; Evans *et al*., 2021), and because the spleen can be an important source or regulator of circulating cytokines (Willis *et al*., 2017; Fonseca *et al*., 2021), we evaluated whether COX-1 inhibition by SC-560 affected the plasma concentrations of the early pro-inflammatory cytokines TNF, IL1β and IL6, as well as the plasma concentration of the anti-inflammatory cytokine IL10. Of these, only TNF and IL1β were detectable in the plasma of the LPS-challenged rats at the time corresponding to the maxima of the cardiovascular, thermal and metabolic responses (80 min post-LPS). SC-560 exerted no effect whatsoever on the concentrations of TNF and IL1β (Fig. 1G-H). Hence, it looks as though cytokine-independent mechanisms are at play when it comes to the role of COX-1 in the severity triad of physiological alterations.

Lastly, a targeted lipidomic approach was used to screen for the COX-1-derived product that plays a role in the early phase of endotoxic shock. Eight products from this pathway were analysed (Fig. 2A). They were not detectable in the plasma of rats not challenged with LPS but became detectable at 80 min post-LPS. Contrary to our expectation, though, SC-560 did not show preference to inhibit a product over others in this phase. On the contrary, except for the stable metabolite of PGI_2_ (PGF_2α_), all products analysed were significantly reduced in the plasma of the LPS-challenged rats pretreated with SC-560 (Fig. 2B). A similar nonselective pattern was observed for the effects of SC-560 on prostanoid concentrations in spleen homogenates (Fig. 2C). We then took a different screening approach, which was based on the premise that if the spleen is a relevant source of the COX-1-derived product that plays a role in the physiological manifestations of severity, the most relevant COX-1-derived product in this context should be changed to equivalent extents in the spleen and plasma. This criterion was best met by PGE_2_, its more stable metabolite (d,k-PGE_2_) and PGD_2_ (Figs. 2D-E). Therefore, PGE_2_ and PGD_2_ should be looked upon as putative candidates for the COX-1-derived products that play roles in the physiological manifestations of severe systemic inflammation.

**Figure 2.**
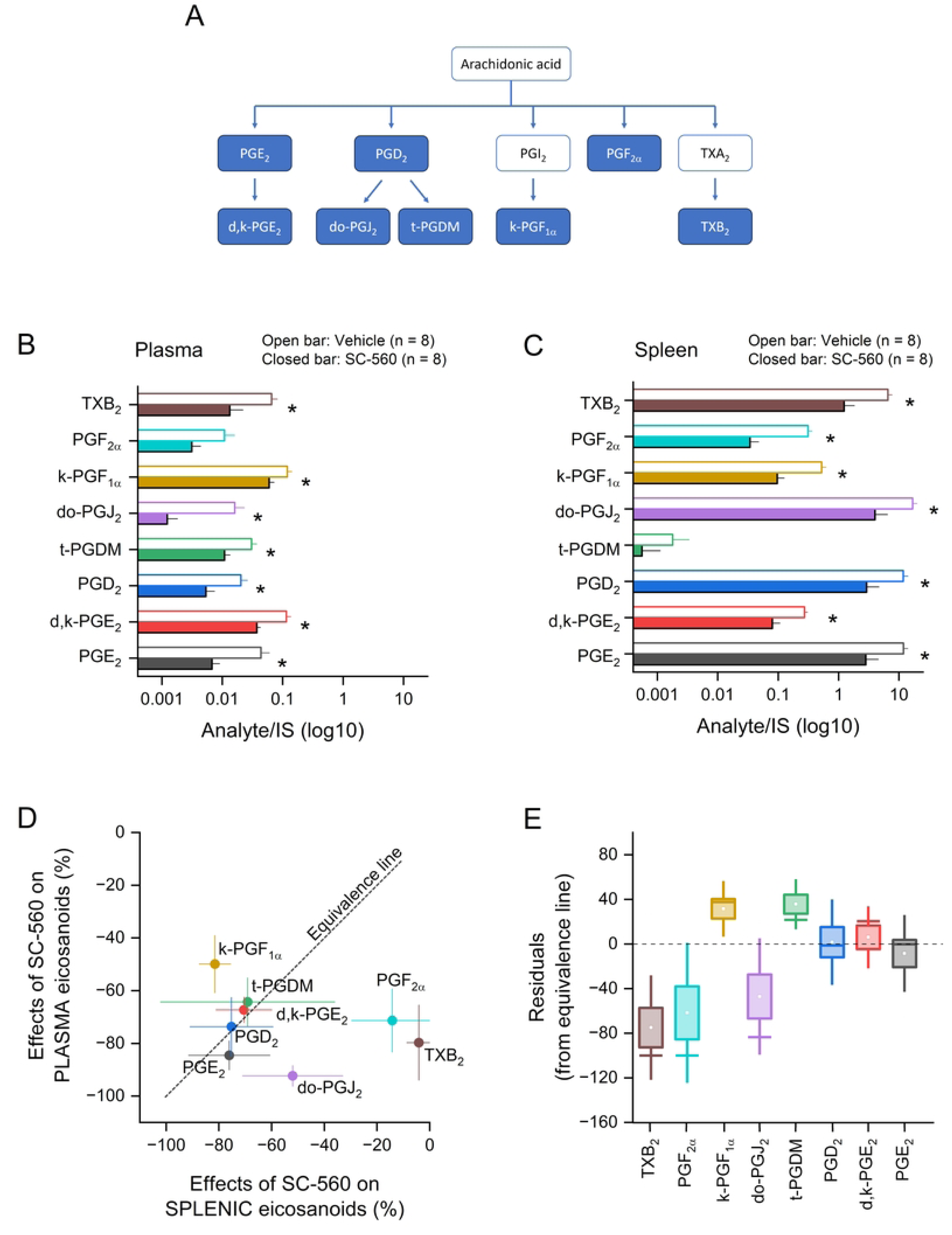
Effects of SC-560 on prostanoid profiles in the plasma and spleen. Panel **A** provides a schematic representation of the eicosanoids and their metabolites that were measured in the LC-MS/MS-based lipidomic analysis. Plasma concentrations and splenic concentrations of eicosanoids at 80 min post-LPS are shown in **B** and **C**, respectively. In panel **D**, the SC-560-induced changes in plasma eicosanoids are plotted against the SC-560-induced changes in splenic eicosanoids. The theoretical equivalence line denotes the points at which plasma and spleen concentrations would have changed to the same extent. Panel **E** depicts how distant (residual) each analyte was from the equivalence line. Data are presented as mean ± SEM. The number of animals per group (n) is indicated. *Statistically significant difference between the SC-560 and vehicle groups.

## 4. Discussion

The present study advances knowledge regarding the role of the constitutive COX isoform (COX-1) in severe cases of acute systemic inflammation. In the rat model of LPS-induced systemic inflammation, we found that COX-1 plays a broad role in several aspects of severe systemic inflammation, namely, the triad of physiological manifestations that accompany it: hypotension, hypothermia and metabolic acidosis. Most importantly, the present study provides direct evidence in support of the hypothesis that the COX-1-dependent eicosanoid storm that drives the severity triad has its origin in the spleen. And it shows that this mechanism is independent of the cytokines typically associated with the early phase of systemic inflammation.

Of the most abundant cellular populations in the spleen, macrophages (Shay *et al*., 2017), T lymphocytes (Pablos *et al*., 1999; Paccani *et al*., 2002) and B lymphocytes (Blaho *et al*., 2009; Baldari, 2014) are known to express functional COX-1. Of these, macrophages are likely to be particularly important in the context of the present study for being activated within minutes of an LPS challenge (Perlik *et al*., 2005; Steiner *et al*., 2006). Platelets are another putative source of eicosanoids in the spleen. One-third of the body’s platelets are known to reside in the spleen (Chamberlain *et al*., 1990), and platelets are renowned for their ability to produce and secrete eicosanoids in a COX-1-dependent fashion (Patrono *et al*., 2001). Moreover, besides serving as a dynamic source of circulating platelets, the splenic platelet pool may also perform spleen-specific functions (Kuwana *et al*., 2002; Horioka *et al*., 2021), so that they can contribute to the eicosanoid storm without leaving the spleen. One interesting mechanistic aspect is that eicosanoid production by macrophages and platelets seem to result not from sensing of LPS itself, but rather from sensing of substances that are released in response to LPS, such as inflammasome-activating substances (von Moltke *et al*., 2012) and complement fragments (Perlik *et al*., 2005). Insofar as these secondary signals reflect the intensity of the body’s response to LPS, this mechanistic framework fits well with the present data implicating COX-1-derived prostanoids in physiological manifestations of severity.

Identifying the COX-1-derived prostanoid(s) that mediate(s) hypotension, hypothermia and acidosis has proven to be a difficult task, because our initial lipidomic screening revealed that most of the prostanoids measured were suppressed by COX-1 inhibition in the plasma or spleen at 80 min post-LPS. We then had to take an alternative, less straightforward approach that was based on the premise that if the spleen is a relevant source of the COX-1-derived product that plays a role in the severity triad, the most relevant COX-1-derived product in this context should be changed to equivalent extents in the spleen and plasma. This strategy appointed at PGE_2_ and PGD_2_ as putative candidates. The candidacy of PGE₂ is highly plausible given its well-established role as a potent vasodilator (Tang *et al*., 2000; Hristovska *et al*., 2007). What is more, a recent study has provided evidence that the early vasodilation that drives LPS-induced hypotension results from heightened autoregulation of blood flow (Moretti *et al*., 2023), a process in which PGE_2_ has been implicated (Vikse & Kiil, 1985; Li *et al*., 1997). Regarding LPS-induced hypothermia, PGD_2_ is the only prostanoid that has been shown to elicit hypometabolism-driven decreases in T_c_ (Ueno *et al*., 1982; Kandasamy & Hunt, 1990), although there is some controversy regarding this effect (Krueger *et al*., 1992; Gao *et al*., 2009a; Gao *et al*., 2009b). Less is known on the relationship of PGE_2_ and PGD_2_ with metabolic acidosis, but two aspects deserve comment. The first aspect is that, in some contexts, PGE_2_ may contribute to sympathetic activation (Ootsuka *et al*., 2008; Rathner *et al*., 2008; Zhang *et al*., 2011) and, as such, it could contribute to the sympathetic activation of glycolysis that mediates, at least partly, acidosis in systemic inflammation (Levy *et al*., 2008). The second point is that acidosis has been shown to be mechanistically independent of hypometabolism/hypothermia in rats challenged with LPS (Steiner *et al*., 2017), so whether or not PDG_2_ mediates hypothermia, it can have a distinct or no effect on the concomitant acidosis. Further research will be needed to test, in direct experiments, the involvement of PGE_2_ and PGD_2_ in the triad of physiological manifestations of systemic inflammation. But, regardless, we think the present findings are of value for setting the foundation for such studies.

In conclusion, the present study establishes the spleen as a critical source of COX-1-derived eicosanoids that mediate the early severity triad (hypotension, hypothermia and acidosis) in the LPS model of systemic inflammation. It also indicates that eicosanoids produced via COX-1 exert these effects without interfering with the cytokine network. Furthermore, our integrated approach identifies PGE_2_ and PGD_2_ as putative candidates for the COX-1-derived products that mediate the severity triad. A better understanding of the mechanisms linking COX-1 and severe systemic inflammation may open avenues for novel therapeutic approaches in severe systemic inflammatory conditions such as sepsis, trauma, cholestasis and pancreatitis.

## Acknowledgments

We thank Mrs. Silvana Silva for technical assistance.

## Ethics Statement

All animal procedures were performed in accordance with institutional requirements and were approved by the Animal Ethics Committee of Universidade de Sao Paulo (approval numbers 015/2015 and 7462060824). The study was also carried out in line with the ARRIVE guidelines.

## Data Availability Statement

The data that support the findings of this study are available from the corresponding author upon reasonable request.

